# Significant expression of FURIN and ACE2 on oral epithelial cells may facilitate the efficiency of SARS-CoV-2 entry

**DOI:** 10.1101/2020.04.18.047951

**Authors:** Mei Zhong, Bing-peng Lin, Hong-bin Gao, Andrew J Young, Xin-hong Wang, Chang Liu, Kai-bin Wu, Ming-xiao Liu, Jian-ming Chen, Jiang-yong Huang, Learn-han Lee, Cui-ling Qi, Lin-hu Ge, Li-jing Wang

## Abstract

**Background:** Leading to a sustained epidemic spread with >2,000,000 confirmed human infections, including >100,000 deaths, COVID-19 was caused by SARS-CoV-2 and resulted in acute respiratory distress syndrome (ARDS) and sepsis, which brought more challenges to the patient’s treatment. The S-glycoprotein, which recognized as the key factor for the entry of SARS-CoV-2 into the cell, contains two functional domains: an ACE2 receptor binding domain and a second domain necessary for fusion of the coronavirus and cell membranes. FURIN activity, exposes the binding and fusion domains, is essential for the zoonotic transmission of SARS-CoV-2. Moreover, it has been reported that ACE2 is likely to be the receptor for SARS-CoV-2. In addition, FURIN enzyme and ACE2 receptor were expressed in airway epithelia, cardiac tissue, and enteric canals, which considered as the potential target organ of the virus. However, report about the expression of FURIN and ACE2 in oral tissues was limited.

**Methods:** In order to investigate the potential infective channel of new coronavirus in oral cavity, we analyze the expression of ACE2 and FURIN that mediate the new coronavirus entry into host cells in oral mucosa using the public single-cell sequence datasets. Furthermore, immunohistochemical staining experiment was performed to confirm the expression of ACE2 and FURIN in the protein level.

**Results:** The bioinformatics results indicated the differential expression of ACE2 and FURIN on epithelial cells of different oral mucosal tissues and the proportion of FURIN-positive cells was obviously higher than that of ACE2-positive cells. IHC experiments revealed that both the ACE2-positive and FURIN-positive cells in the target tissues were mainly positioned in the epithelial layers, partly expressed in fibroblasts, which further confirm the bioinformatics results.

**Conclusions:** Based on these findings, we speculated that SARS-CoV-2 could effectively invade oral mucosal cells though two possible routes: binding to the ACE2 receptor and fusion with cell membrane activated by FURIN protease. Our results indicated that oral mucosa tissues are susceptible to SARS-CoV-2, which provides valuable information for virus-prevention strategy in clinical care as well as daily life.

## 1. Introduction

In December 2019, an increasing number of patients with pneumonia caused by unknown pathogen emerged in Wuhan, Hubei, China, with clinical presentations greatly resembling viral pneumonia ^[1, 2, 3]^. A novel coronavirus, which was named as Corona Virus Disease 19 (COVID-19) by The World Health Organization (WHO), was detected from lower respiratory tract samples via deep sequencing analysis ^[1]^. As of 15th April 2020, the number of coronavirus infections approaches more than two million people and kill more than a hundred thousand ^[4]^. The present outbreak of the 2019 coronavirus strain has been labeled as a global pandemic by WHO ^[5]^. The Covid-19 outbreak has posed critical challenges for the public health, research, and medical communities.

Similar to those of patients with SARS and MERS, the clinical manifestations of COVID-19 at illness onset are fever, dry cough, and myalgia. The patients may suffer from dyspnea, shortness of breath, and then respiratory failure and even death in later stages ^[2, 6, 7]^. Previous findings indicated that the ACE2 plays an important role in cellular entry, thus ACE2-expressing cells may act as target cells and are susceptible to COVID-19 infection ^[2, 8, 9, 10,]^. Besides, ACE2 receptor was proven to be expressed in 72 human tissues, and is abundantly present in the epithelia of the lung and small intestines ^[11]^. Studies also found ACE2 expression in the epithelial layer of nasal and oral mucosa. However, this conclusion was based on the bulk RNA datasets. The cell type and number of cells expressing ACE2 in each tissue could not be estimated.

As a co-expressed membrane endopeptidase of ACE2 receptor, FURIN play a central role in the human immunodeficiency virus type-1 (HIV-1) entering into the host cells through cleaving its envelope glycoprotein ^[12]^. Genomic characterization of the SARS-CoV-2 (severe acute respiratory syndrome coronavirus 2) have revealed that FURIN enzyme could activate the specific site of its spike protein ^[13]^. Meanwhile, Li H. et. al revealed that a putative FURIN-cleavage site reported recently in the spike protein of SARS-CoV-2 may facilitate the virus-cell fusion, which can explain the virus transmission features^[14]^. In addition, Ma Y and co-authors found that presence of FURIN cleavage site, which is absent in other coronavirus, during the course of SARS-CoV-2 infection may cause some distinct clinical symptoms from SARS-CoV infection^[15]^. Similar results were also reported in Li X.’s study from China^[16]^. Hence, the expression and distribution of ACE2 and FURIN in human body were the key factors for the coronavirus to invade host cells. Furthermore, online single-cell sequence datasets analysis unveiled that FURIN was expressed in coronavirus’s potential target organs, such as lung, heart, nose, rectum, colon, intestine and ileum ^[14]^. However, information on the expression and distribution of ACE2 and FURIN in oral cavity were limited.

Single-cell RNA sequencing (scRNA-Seq) technique, which provides an avenue to understand gene regulation networks, metastasis and the complexity of intratumoral cell-to-cell heterogeneity at the single-cell level, has diverse applications on bioinformatics, stem cell differentiation, organ development, whole-tissue subtyping and tumor biology ^[17,18,19]^. It captures cellular differences underlying tumor biology at a higher resolution than regular bulk RNA-seq and has revolutionized studies of cancer mechanisms and personalized medicine ^[20]^. For better understanding the potential infection risk in oral cavity, we explored whether the ACE2 and FURIN were expressed and their expressing cells composition and proportion in different oral tissues based on the public scRNA-seq profiles from GEO public databases. Then IHC analysis was used to further confirm the results in the protein level. These results will provide a theoretical basis for clarifying the possible invading path and damage of SARS-CoV-2 in oral tissues.

## 2. Materials and methods

### 2.1 Public scRNA-seq dataset acquisition

Single-cell RNA-seq dataset(GSE103322) including the gene expression and cell type annotation from human oral squamous cell carcinoma tissue (OSCC) were downloaded from the GEO database (https://www.ncbi.nlm.nih.gov/geo/).

### 2.2 Single-cell sequencing analysis

Single-cell RNA-seq datasets (GSE103322) were downloaded from the GEO database. Datasets of five patients, which have the closest histological features with normal tissue, were selected for futher analysis. The scRNA-seq data sets of GSE103322_HN_UMI_table.txt was used to subsequently analyze based on R package Seurat (Version 3.6.2). To control quality, we removed cells with < 50 genes, and as well as the cells with mitochondrial content higher than 5%. Besides, the genes detected in < 3 cells were filtered out in the function CreateSeuratObject. Subsequently, the data was log-normalized using the function NormalizeData with the default parameters. The FindVariableGenes function was used to determine the highly expressed variable genes, followed by PCA dimensionality reduction by RunPCA function. PCA components, which P value was less than 0.05, were selected to further analyze using Two-dimensional t-distributed stochastic neighbor embedding (tSNE). The setting of k.param was 20 in the function FindNeighbors, and the setting of the resolution was 0.5 in the function FindClusters. Finally, the cell types were assigned based on their canonical markers. Gene expression of different cell types was displayed by the functions of DotPlot and VlnPlot. The significant level was set as 0.05.

### 2.3 Human oral tissue specimens

Twelve normal tissues of oral mucosa (three buccal and gingival tissues, two lip, tongue and palatal tissues respectively) were taken from 12 different patients with clinically diagnosed fibrous epithelial polyp or benign tumor ^[11,21]^, whose average age was 53. Written informed consent was obtained for each participant, and the study was in accordance with the Declaration of Helsinki and the guidelines approved by the Clinical Ethics Committee of Stomatology Hospital of Guangzhou Medical University. Tissue morphology was evaluated in haematoxylin and eosin-stained sections by a qualified pathologist.

### 2.4 Immunohistochemical staining

The selected tissues were fixed in 10% neutral formalin solution and embedded in paraffin. The paraffin sections (4 µm in thickness) were deparaffinized in xylene, rehydrated in a graded series of alcohol, then incubated in citrate buffer (pH 6.0) for 5 min at 120°C, and the endogenous peroxidase was blocked by 0.3% H2O2 for 10 min. Subsequently, they were dried and incubated anti-ACE2 (cat. no. ab15348, Abcam, Cambridge, CB, UK) antibody overnight at 4 °C. The next day, the HRP-conjugated secondary antibody was added to the sections. Peroxidase activity was developed by using DAB for 8 min and counterstained with hematoxylin. The expression levels of ACE2 were evaluated by two experimenters. A qualified pathologist analyzed the staining for structures positive for ACE2. The stain intensity of ACE2 and FURIN was semiquantitatively graded according to the previous methods^[22]^.

### 2.5 Statistical analysis

Statistical significance was determined with a one-way ANOVA followed by Bonferroni’s post-hoc test for multiple group comparisons or Student’s t-test for two group comparisons. For all tests, p < 0.05 or < 0.01 was considered statistically significant or very significant.

## 3 Results

### 3.1 Identification of cell types in the oral mucosal tissues

Given that SARS-CoV-2 invade host cells via ACE2 receptors, we detected the ACE2-expressing cell types by analyzing single-cell RNA sequencing data. 1843 individual cells from the oral mucosa of five patients were analyzed. Using the unsupervised graph-based clustering, we found that at least thirteen distinct cell clusters existed in the oral tissues (Fig. 2A), including epithelial cells (clusters 1, 4, 5), fibroblasts (clusters 0, 2, 7), T-cells (clusters 8, 9), B cells (cluster 12). The heatmap of main corresponding cell marker gene expression profiles across the cell types can be found in Fig. 2B. The marker gene of epithelial cells included KRT6B, GJB2, SFN, and KRT14, fibroblasts included THY1, COL1A1, DCN, ACTA2, B cells included SLAMF7, CD19, CD20 and CD79A and T-cells included CD2, CD3, CD45.

**Fig. 1.**
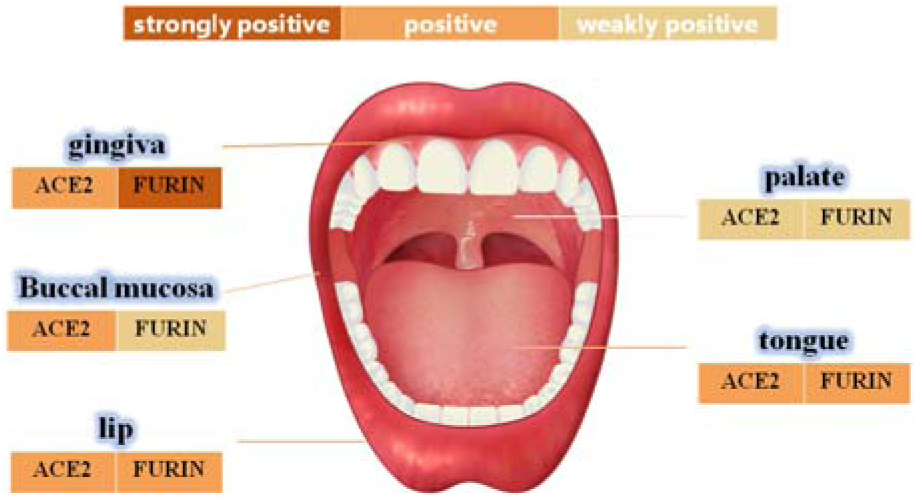
The schematic diagram of FURIN and ACE2 expression levels in different oral sites.

**Fig. 2.**
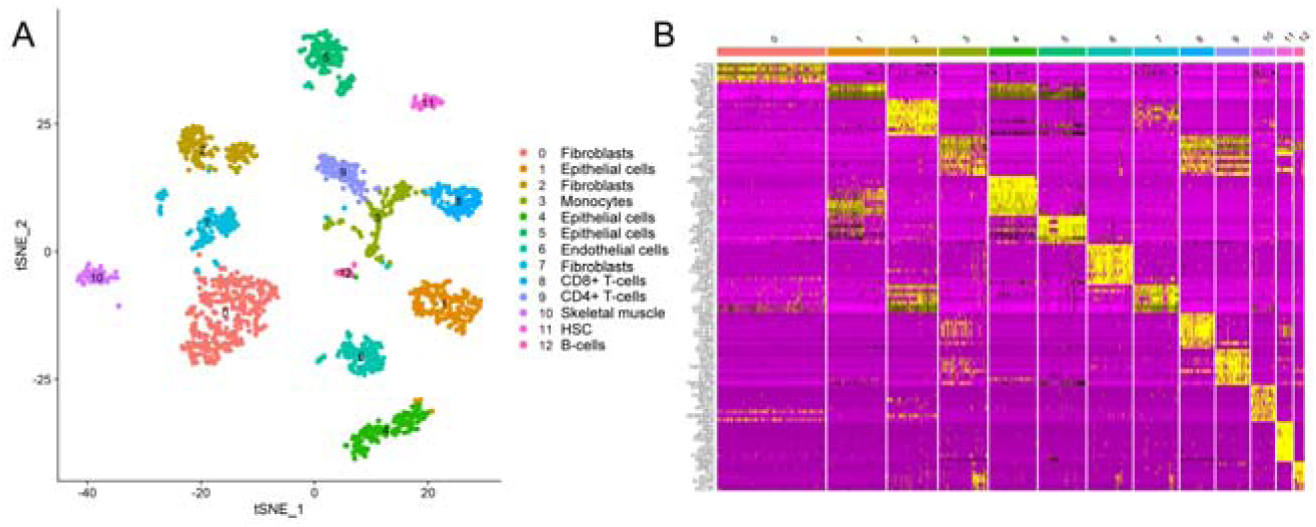
Single-cell atlas of the human oral mucosa. (A and B) Analysis of single-cell sequencing data identified thirteen cell sub-clusters in the oral mucosal cells.

### 3.2 Identification of ACE2-expressing cells in the oral mucosa

As ACE2 had been proven to be one of the major receptors of COVID-2019 in human body, we explored the online datasets to find out the expression level of ACE2 in oral mucosa. As expected, data from GEO indicated that ACE2 receptor expression level was relatively higher in epithelial cells (cluster 1, 4, 5)(Fig. 3A, 3B), while very little cellular expression of ACE2 was found in fibroblasts (cluster 2). Then we calculated the percent of ACE2-expressing cells in the dataset (Fig. 3C). We find that, in oral tissue, the percentage of ACE2-positive cells was 2.2%. Among them, 92% were epithelial cells.

**Fig. 3.**
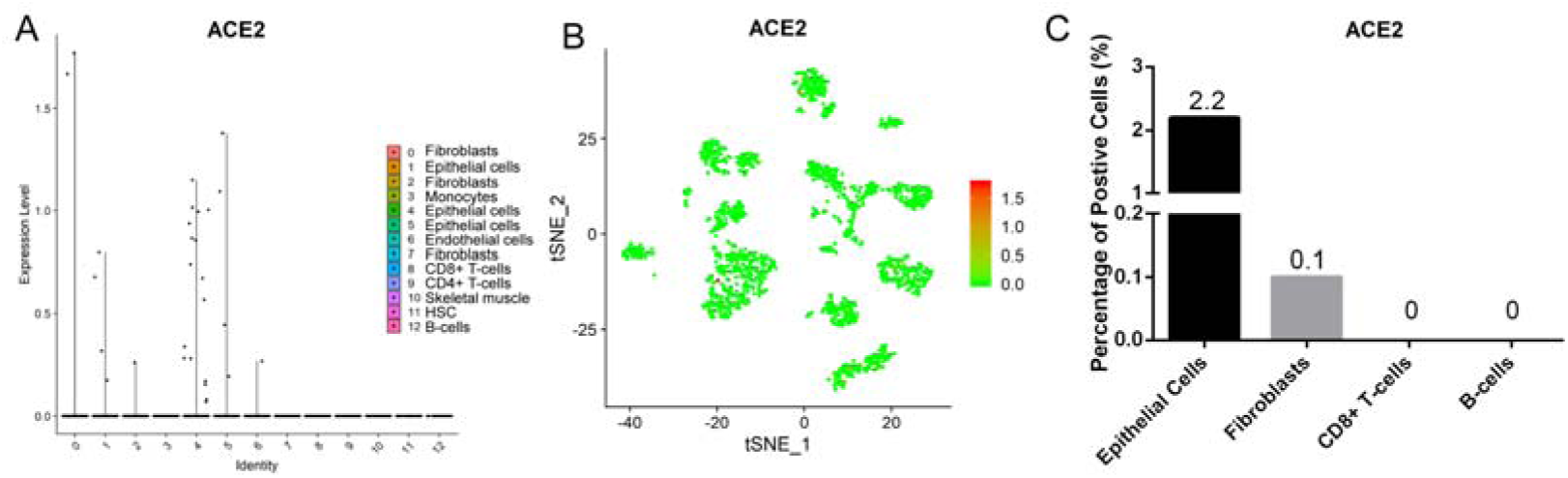
The expression profiles of ACE2 in the oral mucosal cells. (A) Violin and (B) scatter plot showed the expression profile of ACE2 receptor in oral mucosal cells. ACE2 was expressed in epithelial cells. (C) The percentage of ACE2-postive cells in oral mucosa.

### 3.3 Identification of FURIN-expressing cells in the oral mucosa

It has been indicated that FURIN activity is essential for the transmission of SARS-CoV-2. To determine the cell types of expressing FURIN, we analyzed the FURIN mRNA expression using tSNE method. Violin and scatter plot showed that FURIN was highly expressed in epithelial cells, follow by fibroblasts, T-cells, and endothelial cells of the oral mucosal tissues, but hardly expressed in B cells (Fig. 4A, 4B). Meanwhile, the proportion of FURIN-expression cells was approximately 10%. Among them, epithelial cells make up more than 55% (Fig. 4C).

**Fig. 4.**
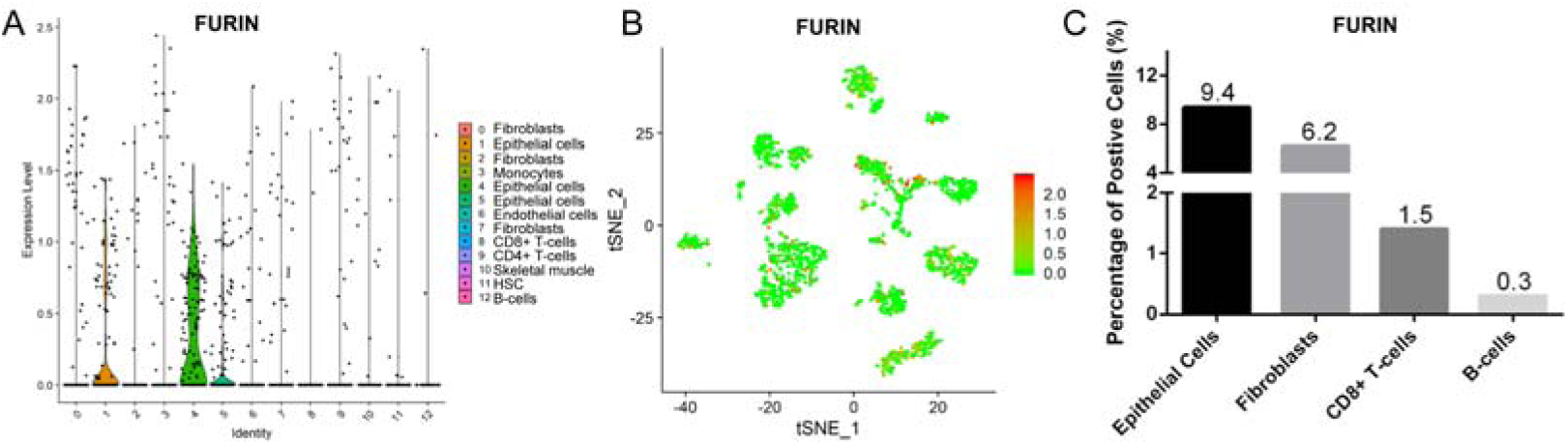
The expression profiles of FURIN in the oral mucosal cells. (A) Violin and (B) scatter plot showed the expression profile of FURIN enzyme in oral mucosal cells. FURIN was highly expressed in epithelial cells. (C) The percentage of FURIN-expressed cells in oral mucosal cells.

### 3.4 ACE2 protein was highly expressed in epithelial layer of the buccal mucosa, lip, palate, tongue and gingiva

To further determine the protein expression level of ACE2 in the oral mucosa, IHC experiments were performed. The first remarkable finding was that ACE2 was present in epithelial cells in all the tissues studied (Fig. 5). Results indicated that the expression level of ACE2 protein was significantly higher in the lip, tongue, and buccal mucosa, especially the epithelial cells in the basal layer, although the mRNA expression level is not as high (Fig. 3). The ACE2-postive cells in the gingival and palate tissue were limited. The corresponding gingiva and palate epithelial cells showed weak positive ACE2 staining. Also, we found that ACE2 expressed in fibroblasts and endothelial cells. These results indicate that oral epithelial cells were the potential targets of SARS-CoV-2.

**Fig. 5.**
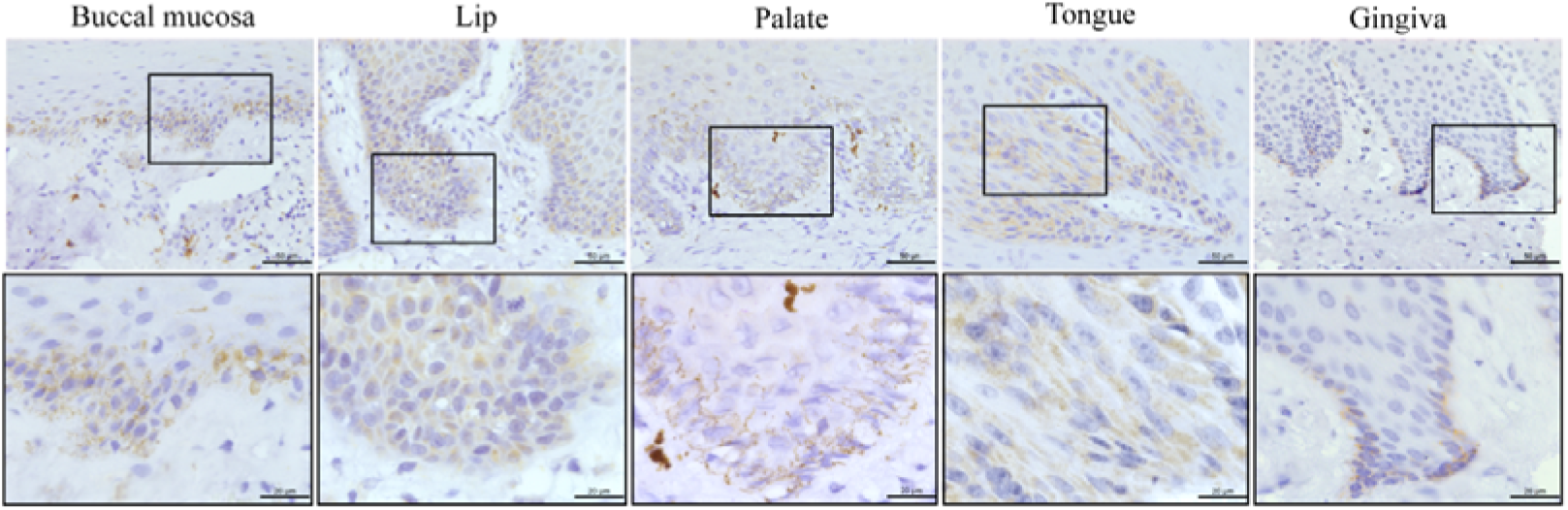
ACE2 protein was highly expressed in the various oral tissues. Immunohistological staining against ACE2 was used to determine whether ACE2 express in the normal buccal, lip, palate, tongue, and gingival tissues from the human. The lower panels demonstrate the appearance of basal layer and epithelial cells at higher magnification as indicated by black squares in upper panels. ACE2 was highly expressed in basal layer epithelial cells of these five oral tissues. Scale bars = 50 μm.

### 3.5 FURIN protein was highly expressed in the normal oral mucosa

FURIN mRNA expression has been described mainly in oral epithelial cells through scRNA-seq technique. To determine whether this differential expression of FURIN was maintained at the protein level, oral tissues from different sites were screened. As shown in Fig. 6, moderate to mild immunostaining in the cytoplasm of epithelial cells can be seen in five normal oral tissues. Surprisingly, except for the positive cells in the basal layer of oral epithelium, the spinous layer in all examined tissues also turned out with large numbers of FURIN-positive cells. In addition, the percentage of FURIN-positive cells in lip, tongue and gingiva was higher than that of buccal and palate mucosa. These results indicated that SARS-CoV-2 might be able to infect oral tissues through the route of FURIN-mediated fusion with oral mucosal cell.

**Fig. 6.**
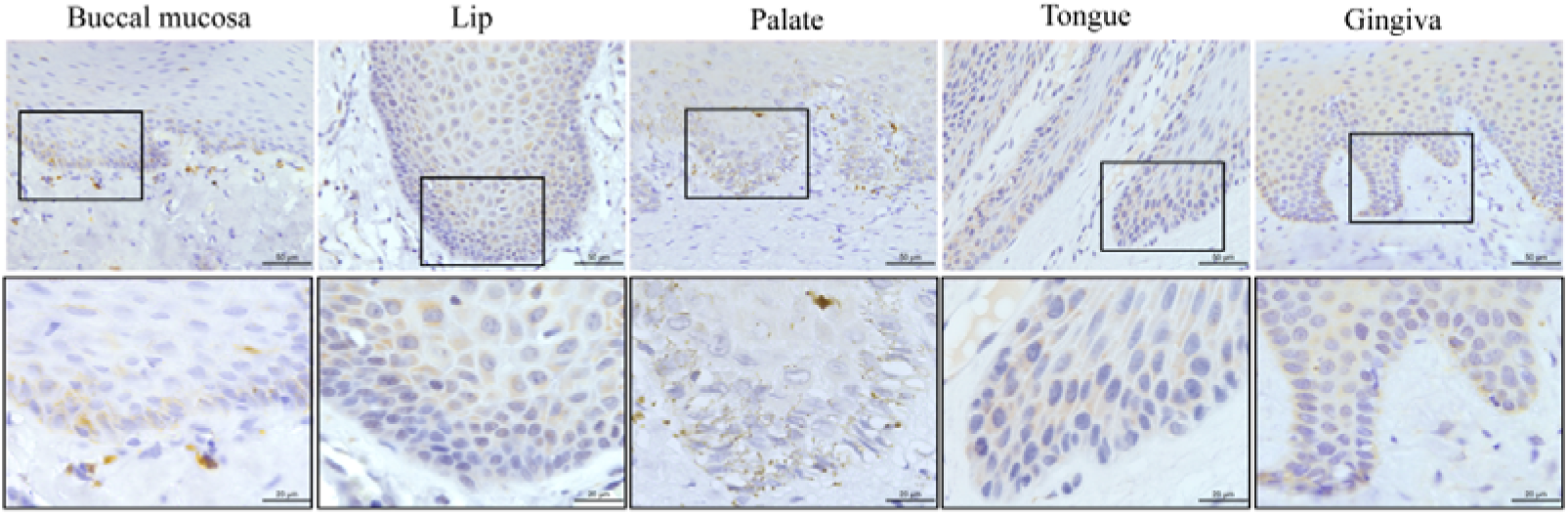
FURIN was highly expressed in the various oral tissues. Immunohistological staining against FURIN was used to determine whether FURIN expressed in the normal buccal mucosa, lip, palate, tongue, and gingival tissues from the patients. The lower panels demonstrate the appearance of basal layer and epithelial cells at higher magnification as indicated by black squares in upper panels. FURIN was highly expressed in spinous and basal layer cells, while FURIN-positive fibroblasts and endothelial cells in tongue, buccal and palate tissues were limited. Scale bars = 50 μm.

## Discussion

The amazing spreading speed of SARS-CoV-2 and the large number of infected people have quickly overwhelmed the public health services in the world, and have become a global public health problem ^[23]^. As mentioned before, ACE2 receptor in cell membrane and FURIN clearage site of SARS-CoV-2 were the key factors that allowed the virus to invade the host ^[14, 15]^. Thus, clarifying the distribution of ACE2 and FURIN at the protein and gene levels in tissues is essential for understanding the pathogenic mechanism of SARS-CoV-2.

Many studies have shown that ACE2 receptor was highly expressed in respiratory epithelium, kidney, cardiovascular system and testis ^[24]^. Some scholars also found that ACE2 was expressed in oral tissues with bulk RNA sequencing data, which could not accurately reflect the expression of ACE2 on the single cell level ^[21, 25]^. Chen et al. benefited from the single-cell sequencing data to further proved the expression of ACE2 receptor in tongue, buccal mucosa and gingiva ^[11]^. However, these studies were based on public databases and were not analyzed together, and further validation of protein levels was lacking. In the current study, we firstly characterized the ACE2 protein expression profiles in five different oral tissues through immunohistochemical staining experiment. Results indicated that ACE2 was mainly present in epithelial cells in all included tissues, especially in buccal mucosa, lip and tongue. Previous study discovered the similar result^[21]^. Overall, these results demonstrated that, compared with other oral sites, buccal mucosa, lip and tongue might be more susceptible to SARS-CoV-2.

In addition, virologist Gary Whittaker of Cornell University, whose team published another structural analysis of the coronavirus S protein on bioRxiv on February 18th, argues that the FURIN cutting site allows the virus to enter cells in a very different way than SARS, and this may affect the stability and spread of the virus ^[26]^. Other research teams have also identified this activation site, suggesting that it may enable the virus to spread effectively between humans^[13]^. It has been demonstrated that SARS-CoV-2 could fuse with cell membrane and then entry into the target cells^[27, 28]^. Researchers have found FURIN proteases in many tissues, including the lungs, liver, and small intestine ^[29]^. However, limited reports were seen regarding the expression of FURIN in oral cavity and whether the new coronavirus could spread by oral cavity are not fully understood. Thus, we investigated the FURIN enzyme in both gene and protein level from five different oral sites and the results would be valuable for the prevention of the COVID-19 spreading.

Previous analysis about the SARS-CoV-2 receptor in the sing cell level were based on the complete OSCC database named GS103322, which was significantly different from the genetic profiles of normal oral tissues^[30]^. In order to accurately analyzing the expression status of FURIN in the gene level, datasets from five OSCC patients, which were graded as 1 and T1N0 stage were selected originated from the online platform in the present research. The obtained single cell RNA-seq data were processed and analyzed with R software. As expected, FURIN were found abundantly enriched in oral mucosa cells, particularly in epithelial cells and the proportion of FURIN-positive cells was higher than that of ACE2-positive cells.

Except for the expression profiles in the gene level, FURIN protein expression in oral cavity was crucial for further clarifying the impact of SARS-CoV-2 on the mouth. Previous reports had demonstrated that FURIN protein was expressed in metastasizing OSCCs^[31]^. Besides, FURIN was expressed in normal epithelium and up-regulated in SCCs from three different organs^[22]^. But the relative report of FURIN protein expression in normal oral cavity was limited. For further validating the results in the gene level, FURIN immunohistochemistry was therefore performed on conventional sections for the first time to assess its expression in oral mucosa obtained from five different sites. It is noteworthy that moderate positive expression of FURIN protein were observed in tongue, giniva and lip, while weakly positive in the buccal and palatal tissues. As a result, we speculated that SARS-CoV-2 could fuse with the FURIN enzyme on the cell membrane the oral mucosal cells, then entry into the host cells. Alternatively, FURIN could be recognised as the potential drug target of suppressing the infection of new coronavirus.

Strikingly, the scRNA-Seq data analysis and IHC tests results indicated that ACE2 and FURIN were both abundantly expressed in oral mucosa, especially on oral epithelial cells, which were similar to the previous reports ^[32, 33, 34]^. In other word, two key proteins of SARS-CoV-2 entering the target cell in different routes were both present in oral mucosa, which implied the potential attack and spread of SARS-CoV-2 in the oral tissues. Moreover, the live SARS-CoV-2 was consistently detected in the self-collected saliva of 91.7% of patients till 11 days after hospitalization, and it could be transmitted via saliva, directly or indirectly, even among asymptomatic infected patients^[35]^. Thus, it is suggested that all the healthcare workers should focus on taking effective measures to prevent the virus from spreading via the oral cavity and potential impairment of oral tissues should be evaluated after infected with SARS-CoV-2, giving more evidence to the prevention and treatment of COVID-19. We should also pay close attention to protecting the oral tissues of all people, preventing the virus from spreading through the mouth.

In the current study, we investigated the expression of other endopeptidases in various oral mucosa cells using the public single-cell sequence datasets. As shown by voilin plot in supplemental files, the high expression of ACE2 sheddase, ADAM17, ADAM10 indicated the higher membrane fusion activity, virus uptake and virus replication in the oral mucosa. Besides, the endopeptidases (such as CAPN1 and CAPNS1) were abundantly enriched in the oral mucosal cells. Large proportion of endopeptidases positive cells in the different oral sites provided opportunities for SARS-CoV-2 infection and rapid replication in the oral cavity ^[15]^. These results were in accordance with the previous study, which indicated the residence of SARS-CoV-2 in nasal and throats swab samples, suggesting the strong spread ability in oral cavity ^[36]^.

In conclusion, our results systematicly investigated the expression profiles of FURIN enzyme and ACE2 receptor in oral cavity at the gene and protein level for the first time. Furthermore, significant expression of FURIN and ACE2 were discovered in oral epithelial cells, implying the possible two transmission routes of the coronavirus through oral mucosa, which provides a new insight to the future prevention strategy in clinical cares as well as daily life. More evidence is still needed to reinforce these findings.

## Declarations

### Ethics approval and consent to participate

All human participants signed the informed consent before collecting the samples. Each of the human studies was reviewed and approved by the Institution Ethics Committee of Stomatology Hospital of Guangzhou Medical University.

### Consent for publication

Not applicable.

### Availability of data and material

The datasets supporting the results of this article are included within the article (and its additional files).

### Authors’ contributions

LBP, ZM and GHB helped in the sample collection, gathering of participant data, writing original draft. AJY helped in the manuscript writing, data analysis, and interpretation. WXH, LC, WKB and LMX helped in the IHC assays, and final approval of manuscript. LLH, QCL, HJY and CJM helped in the research design, analysis and interpretation of results. WLJ and GLH conceived of the study, and participated in its design and coordination. All authors read and approved the final manuscript.

## Acknowledgements

We thank all of the volunteers who agreed to participate in this study, as well as Wang Xinran for his contributions to this work.

## Authors’ information

Not applicable

## Supplemental Files

**Table S1.** Patients and samples included in dataset

| Designation | Age/Sex | Primary site | Pathologic stage | Grade | LVI |
| --- | --- | --- | --- | --- | --- |
| MEP 6 | 88/F | right floor of mouth | T4aN2c | 1 | Absent |
| MEP 8 | 82/F | right hard palate | T4aN0 | 1 | Absent |
| MEP 16 | 63/F | left lateral tongue | T2N0 | 1 | Absent |
| MEP 22 | 77/M | left buccal mucosa | T1N0 | 2 | Absent |
| MEP 28 | 58/M | right lateral tongue | T2N2c | 1 | Present |

**Fig. S2.**
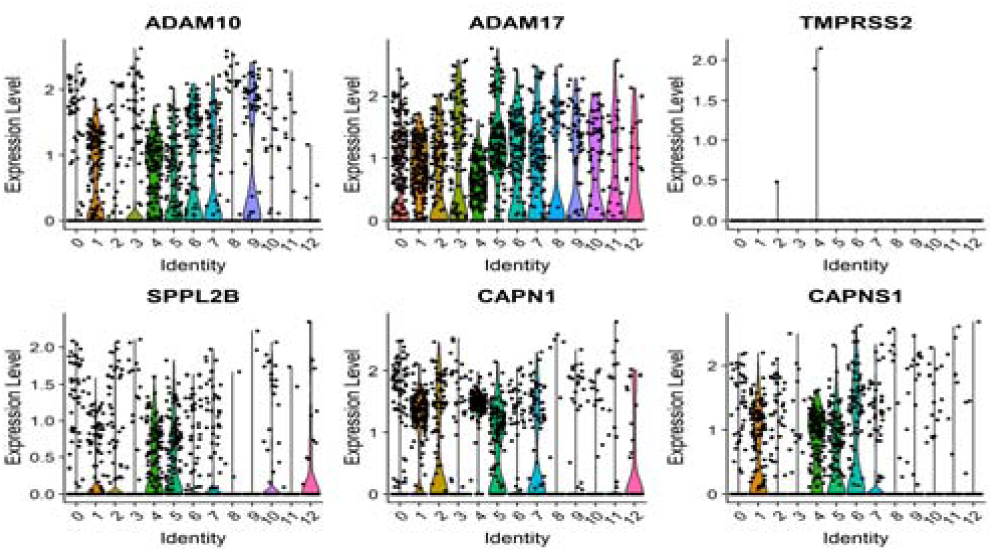
The expression profiles of FURIN in the oral mucosal cells. (A) Violin and (B) scatter plot showed the expression profile of FURIN receptor in the oral mucosal cells. FURIN was highly expressed in epthelial cells. (C) The percentage of FURIN-expression cells in the adult oral mucosal cells.

**Fig. S3.**
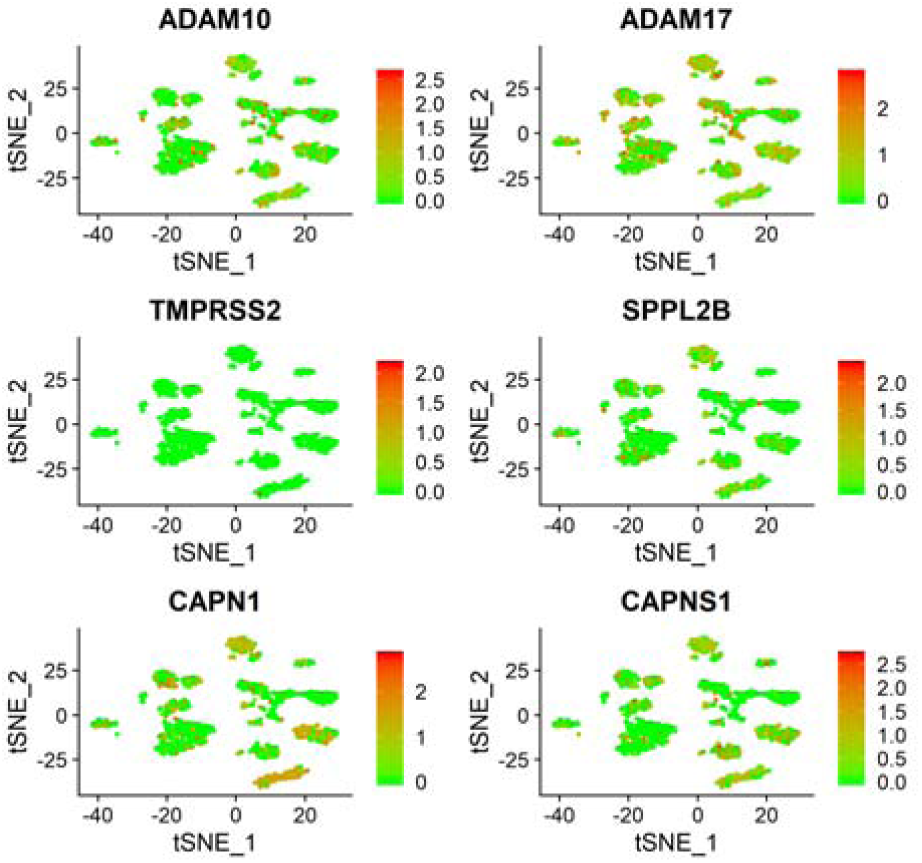
The expression profiles of other endopeptidase in the oral mucosal cells. Scatter plot showed the distribution profile of ADAM10, ADAM17, TMPRSS2, SPPL2B, CAPN1 and CAPNS1 in the oral mucosal cells.

